# Limited contribution of rare, noncoding variation to autism spectrum disorder from sequencing of 2,076 genomes in quartet families

**DOI:** 10.1101/127043

**Authors:** Donna M. Werling, Harrison Brand, Joon-Yong An, Matthew R. Stone, Joseph T. Glessner, Lingxue Zhu, Ryan L. Collins, Shan Dong, Ryan M. Layer, Eirene Markenscoff-Papadimitriou, Andrew Farrell, Grace B. Schwartz, Benjamin B. Currall, Jeanselle Dea, Clif Duhn, Carolyn Erdman, Michael Gilson, Robert E. Handsaker, Seva Kashin, Lambertus Klei, Jeffrey D. Mandell, Tomasz J. Nowakowski, Yuwen Liu, Sirisha Pochareddy, Louw Smith, Michael F. Walker, Harold Z. Wang, Mathew J. Waterman, Xin He, Arnold R. Kriegstein, John L. Rubenstein, Nenad Sestan, Steven A. McCarroll, Ben M. Neale, Hilary Coon, A. Jeremy Willsey, Joseph D. Buxbaum, Mark J. Daly, Matthew W. State, Aaron Quinlan, Gabor T. Marth, Kathryn Roeder, Bernie Devlin, Michael E. Talkowski, Stephan J. Sanders

## Abstract

Genomic studies to date in autism spectrum disorder (ASD) have largely focused on newly arising mutations that disrupt protein coding sequence and strongly influence risk. We evaluate the contribution of noncoding regulatory variation across the size and frequency spectrum through whole genome sequencing of 519 ASD cases, their unaffected sibling controls, and parents. Cases carry a small excess of *de novo* (1.02-fold) noncoding variants, which is not significant after correcting for paternal age. Assessing 51,801 regulatory classes, no category is significantly associated with ASD after correction for multiple testing. The strongest signals are observed in coding regions, including structural variation not detected by previous technologies and missense variation. While rare noncoding variation likely contributes to risk in neurodevelopmental disorders, no category of variation has impact equivalent to loss-of-function mutations. Average effect sizes are likely to be smaller than that for coding variation, requiring substantially larger samples to quantify this risk.

## Introduction

The rapid progression of genomics technologies, coupled with expanding cohort sizes, have led to significant progress in characterizing the genetics of autism spectrum disorder (ASD). To date, studies of ASD cohorts have included genotyping array technologies to survey large copy number variations (CNVs)^1-6^ and common variants,^7,8^ exome sequencing to scan the protein coding genome,^1,9-16^ and long-insert sequencing to identify large chromosomal abnormalities.^17,18^ While genetic variation across the allele frequency spectrum influences ASD risk,^19^ robust discovery of specific genetic loci has been driven by the identification of extremely rare *de novo* mutations that are predicted to disrupt protein coding genes. Since these mutations are newly arising in the child, they receive limited exposure to natural selection and can therefore exert considerable risk for ASD, given the well documented reduction in fecundity in ASD cases.^20^ Two factors have driven locus discovery in ASD: the presence of critical sites in coding genes that, when mutated, severely disrupt gene function leading to dramatic biological consequences, and the ability to predict such disruption based on gene models, either through large-scale deletion or the annotation of point mutations using the triplet genetic code.

Most ASD subjects do not carry either gene disrupting point mutations or large *de novo* CNVs,^1^ hence assaying *de novo* noncoding mutations could identify uncharacterized reservoirs of genetic risk. Yet, while the vast majority of *de novo* mutations (97%) arise outside the coding genome, they present an interpretive challenge. Unlike the coding region, we do not have the same cipher, the triplet code, to predict which nucleotides will critically alter gene function when mutated and which will be functionally inert. Association of noncoding variation with complex traits is well-documented, with the overwhelming majority being common variants mapping outside of gene regions and often in proximity to putative regulatory domains. These common variant associations typically have modest effect sizes. While the impact on gene expression levels, splicing events, or other regulatory processes is defined for some noncoding associations, the key regulatory consequences remain unknown for the majority. This uncertainty in functional prediction necessitates an unbiased approach to rare variant disease association from WGS that parallels the statistical rigor applied to common variant analyses.

In the case of coding variation, the analysis of *de novo* mutations allows identification of an extremely rare class of variation and the ability to unequivocally link it to disease. We hypothesized that if a class of noncoding variation has a similar impact, then analysis of *de novo* mutations presents a powerful approach to discovery. Furthermore, disruption of specific regulatory elements could provide key insights into the cell types, brain regions, and developmental periods critical to neurodevelopmental disorders.^21-23^ The success of this approach will be dependent on the number of critical sites, the susceptibility of these sites to mutations, and our ability to predict disruption of these sites.

Here, we present an exploration of the impact of noncoding regulatory variation from WGS in a cohort of 519 ASD cases, their unaffected siblings as controls, and both parents (2,076 individuals) from the Simons Simplex Collection (SSC).^24^ We find no specific category of *de novo* or rare noncoding regulatory variation that reaches statistical significance when accounting for the tests we performed in this framework. For ASD – and likely other common, complex disorders – these results indicate that there is no known category of functional annotation in the noncoding genome that confers comparable risk to *de novo* loss-of-function coding mutations. Our results underscore the challenges in the analysis of noncoding variation: 1) absence of a noncoding equivalent to the triplet genetic code to determine which *de novo* variants will be functionally relevant and which will be silent; 2) the size of the noncoding genome; and 3) the necessity of testing a multiplicity of hypotheses due to the numerous classes of noncoding functional elements and types of genomic variation. We conclude that the average relative risk (RR) of rare noncoding variants will be modest, they will be distributed widely across the genome, and sample sizes required to identify them will need to be substantially larger.

### Cohort selection and characteristics

All 519 cases were selected from the SSC based on the absence of *de novo* loss-of-function mutations or large *de novo* CNVs in prior data, with the objective of enriching for undiscovered *de novo* variation. The majority of cases (92%, N=480/519) were selected randomly after this exclusion, however the remaining 8% were selected for a pilot study^25^ to increase the representation of older fathers, female cases, and cases with comorbid intellectual disability (ID; defined here as nonverbal IQ ≤70), all of which have been associated with increased rates of protein-damaging mutations.^1^ Of the 519 WGS cases, 10.6% are female, which is lower than the 15.0% (p = 0.02) in cases excluded due to known *de novo* mutations and the 14.1% (p = 0.04) in the remainder of the SSC without WGS data. No significant differences were observed in the fraction of cases with ID, which were 25.8%, 26.0% and 25.2%, respectively.

The contribution of coding *de novo* mutations to neurodevelopmental disorders is a continuum ranging from severe intellectual disability, with *de novo* loss-of-function mutations contributing risk in 18% of cases in the Deciphering Developmental Disorders (DDD) cohort,^26^ to later-onset disorders, such as schizophrenia in which *de novo* loss-of-function mutations are unlikely to contribute to more than 2% of cases. ASD falls between these two extremes, with about 7% of SSC cases carrying such mutations. The contribution of inherited (largely common) variation appears to run in the opposite direction, as reflected by the high sibling recurrence rates in ASD^27^ and schizophrenia^28^ compared to ID cases.^29^ Given this relationship, we predicted common variant ASD burden from microarray data of the 1,631 families in the SSC of European ancestry (Extended Data Fig. 1). As expected, we observed a lower burden of common variant risk in cases excluded due to known *de novo* mutations than in our WGS cohort and the remainder of the SSC (p=0.03, one-sided t-test), but no difference between our cohort and the remainder of the SSC.

### Single nucleotide variants and insertion-deletions

Single nucleotide variants (SNVs) and small insertion-deletions <50 bp (indels) were discovered in the new WGS subset using the Genome Analysis ToolKit (GATK),^30^ and family structure was leveraged to define high quality calls (Extended Data Fig. 2-5). Overall, we identified 3.7 million high quality, autosomal variants per individual, including 3.4 million SNVs and 0.3 million indels. From these variants, *de novo* SNVs and indels were predicted using multiple detection algorithms and excluding low complexity regions. These predictions were ensured to be of high confidence by tuning and subsequent validation (Extended Data Fig. 5-6). Confirmation rates compared favorably with published literature for both SNVs (96.8%, 212/219) and indels (82.4%, 145/176).^25^ Both WGS and whole exome sequencing (WES) data were available for 991 children. Within Gencode-defined, autosomal coding regions, 1,071 *de novo* SNVs and 41 *de novo* indels were detected by WGS compared to 869 *de novo* SNVs and 27 *de novo* indels by WES. Of the 896 *de novo* WES variants, 870 were detected by GATK in the WGS data (97%; 849 SNVs, 21 indels) and 768 of these variants met our high quality *de novo* criteria (88% of 870; 754 SNVs, 14 indels). WGS identified an additional 344 high quality *de novo* mutations (317 SNVs, 27 indels) that were not reported by WES, in large part due to limited coverage in the WES data.

In WGS data we observed a median of 64 *de novo* SNVs and 5 *de novo* indels per child, with a slight excess of mutations in cases compared to their sibling controls after adjusting for quality metrics influencing *de novo* mutation detection using linear regression (RR = 1.024, p = 0.002 for all variants; RR = 1.023, p = 0.003 for noncoding mutations alone). However, when we correct for the effect of paternal age, which is known to affect mutation rates,^10,31^ no significant difference in *de novo* burden remained for all mutations (RR = 1.008, p = 0.28; Extended Data Fig. 8) or noncoding mutations alone (RR = 1.007, p = 0.33). The slight excess of about one noncoding mutation per case, prior to adjusting for paternal age, is likely due to the fact that 56% of cases were born after their sibling controls. This bias towards later born cases is consistent with a wide range of scenarios, only one of which involves a direct relationship between noncoding *de novo* mutation and ASD risk. Regardless of the mechanism, this modest excess in cases will confound a search for the noncoding elements that mediate ASD risk, therefore, correction for all covariates, including paternal age, was applied to all subsequent tests of *de novo* burden.

The sheer diversity and complexity of noncoding functional annotations necessitates a strategy to interpret the multiple parallel hypotheses. We first assessed whether there was evidence of an excess of variants in cases within regions of the genome defined by genes. As noted, the cohort included only cases that did not carry a *de novo* loss-of-function coding mutation in prior analyses by WES.^1^ Using Gencode gene definitions, we surveyed four coding categories, e. g. missense, and seven noncoding categories, e. g. UTRs (Fig. 1). In all analyses, we tested for an enrichment of mutations mapping to these regions in cases compared to their sibling controls, and then assessed the significance of this enrichment using 10,000 case/control label-swapping permutations comparing the number of *de novo* mutations corrected for paternal age and sequencing quality metrics. This analytical approach is used throughout the manuscript, unless otherwise noted. After correcting for multiple comparisons, no significant excess of *de novo* variants in any gene-defined category was observed. We repeated the analysis considering SNVs and indels separately (Extended Data Fig. 9-10), and considering only variants within or near to one of 179 genes associated with ASD at a liberally defined false discovery rate (FDR < 0.3).^1^ Only an excess of *de novo* missense mutations is apparent (Fig. 1), though both promoter regions and UTRs showed a trend towards enrichment in cases. Substituting ASD-associated genes for constrained genes^32^ or mRNA targets of Fragile X Mental Retardation Protein (FMRP)^33^ did not yield any nominally significant categories, including missense variants, nor did considering only variants at nucleotides conserved across species.

**Figure 1.**
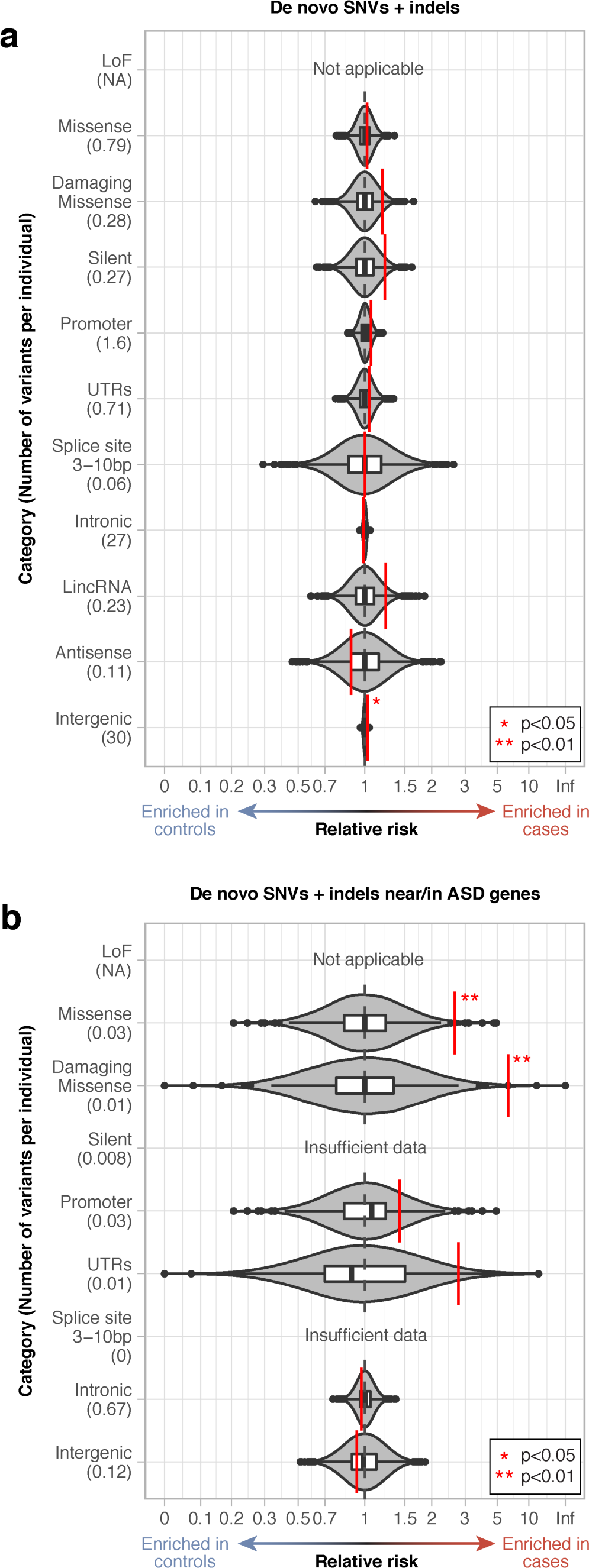
Burden analyses for gene-defined annotation categories. ***a)*** The observed relative risk of de novo mutations in cases vs. controls is shown by the red line against grey violin plots representing 10,000 label-swapping permutations of case-control status for 11 gene-defined annotation categories. Uncorrected p-values are highlighted with red asterisks; the absence of an asterisk indicates the category did not reach nominal significance. Loss-of-function variants were not analyzed as cases with such mutations were excluded from the cohort. **b)** The analysis in ‘a’ is repeated considering only de novo mutations in or near 179 ASD genes.

We next designed an unbiased WGS-association framework for the noncoding genome in ASD. We integrated five approaches to annotation: 1) ASD-associated gene lists (e. g., targets of FMRP); 2) functional annotation (e. g., chromatin state); 3) conservation across species; 4) type of variant (SNVs, indel); and 5) gene-defined categories described above. In total we surveyed 51,801 non-redundant annotation categories derived from combinations of these five annotation approaches. In the absence of a clear *a priori* hypothesis, we treated all of these category comparisons equally and compared the burden of *de novo* mutations in cases vs. controls (Fig. 2a) in a category-wide association study (CWAS). The most strongly associated categories were from coding variants, while the top noncoding category was from mutations underlying H3K36me3 peaks that were nearer to lincRNAs than to other transcripts (Table 1).

**Figure 2.**
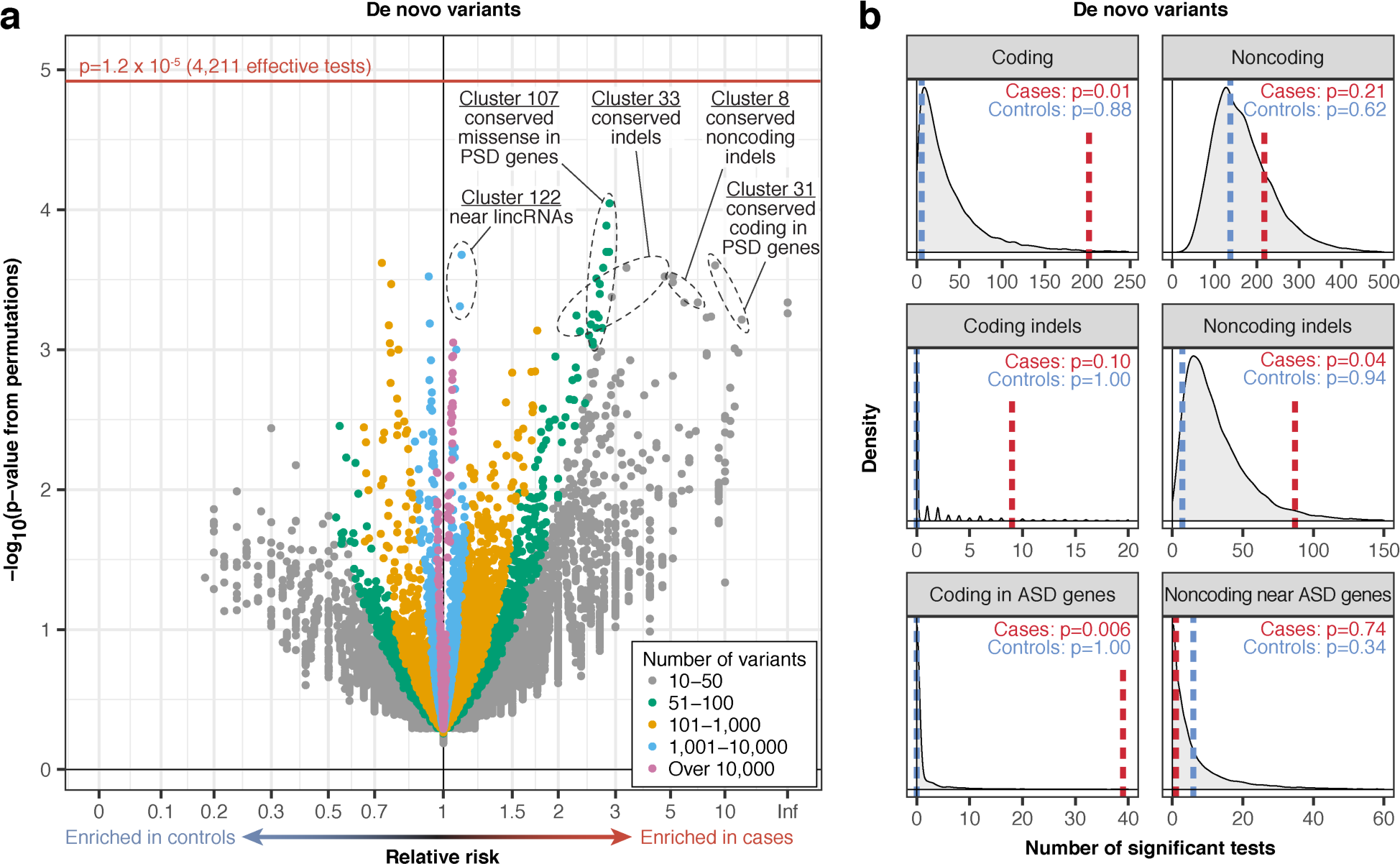
Category-wide association study. **a)** The burden of de novo mutations in cases vs. controls was tested for 51,801 annotation categories. The 11,876 categories with ≥10 observed variants are shown as points in the volcano plot colored by the number of observed mutations. P-values were calculated by 10,000 label-swapping permutations of case-control status in each annotation category. No test exceeds the correction for 4,211 effective tests (horizontal red line). **b)** The number of nominally significant annotation categories (p≤0.05) was calculated for cases (red line), controls (blue line), and 10,000 permutations (grey density plot) to assess whether more annotation categories are enriched for de novo variants in cases than expected in ‘a’. Cases have a greater than expected number of nominally significant categories relating to coding mutations and noncoding indels, but no to all noncoding mutations. P-values were calculated by comparison to the permutation results.

**Table 1.**
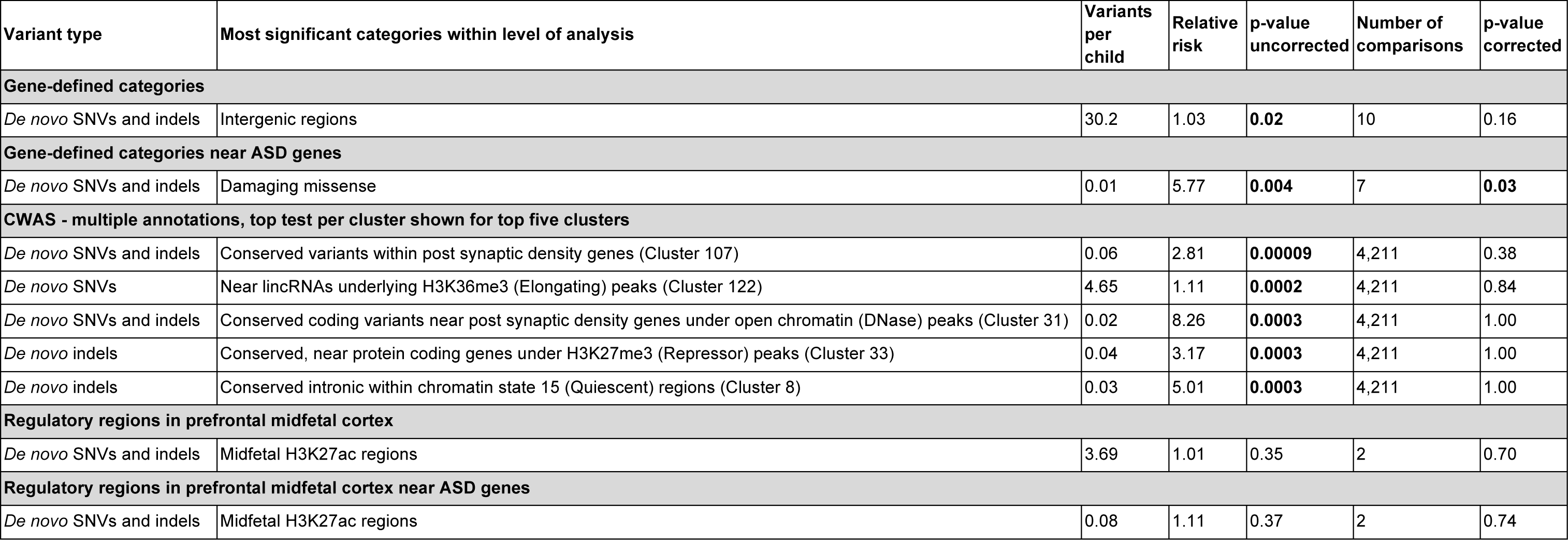
Burden results for most significant or previously implicated annotation categories

| Variant type | Most significant categories within level of analysis | Variants per child | Relative risk | p-value uncorrected | Number of comparisons | p-value corrected |
| --- | --- | --- | --- | --- | --- | --- |
| **Gene-defined categories** |  |  |  |  |  |  |
| *De novo* SNVs and indels | Intergenic regions | 30.2 | 1.03 | **0.02** | 10 | 0.16 |
| **Gene-defined categories near ASD genes** |  |  |  |  |  |  |
| *De novo* SNVs and indels | Damaging missense | 0.01 | 5.77 | **0.004** | 7 | **0.03** |
| **CWAS - multiple annotations, top test per cluster shown for top five clusters** |  |  |  |  |  |  |
| *De novo* SNVs and indels | Conserved variants within post synaptic density genes (Cluster 107) | 0.06 | 2.81 | **0.00009** | 4,211 | 0.38 |
| *De novo* SNVs | Near lincRNAs underlying H3K36me3 (Elongating) peaks (Cluster 122) | 4.65 | 1.11 | **0.0002** | 4,211 | 0.84 |
| *De novo* SNVs and indels | Conserved coding variants near post synaptic density genes under open chromatin (DNase) peaks (Cluster 31) | 0.02 | 8.26 | **0.0003** | 4,211 | 1.00 |
| *De novo* indels | Conserved, near protein coding genes under H3K27me3 (Repressor) peaks (Cluster 33) | 0.04 | 3.17 | **0.0003** | 4,211 | 1.00 |
| *De novo* indels | Conserved intronic within chromatin state 15 (Quiescent) regions (Cluster 8) | 0.03 | 5.01 | **0.0003** | 4,211 | 1.00 |
| **Regulatory regions in prefrontal midfetal cortex** |  |  |  |  |  |  |
| *De novo* SNVs and indels | Midfetal H3K27ac regions | 3.69 | 1.01 | 0.35 | 2 | 0.70 |
| **Regulatory regions in prefrontal midfetal cortex near ASD genes** |  |  |  |  |  |  |
| *De novo* SNVs and indels | Midfetal H3K27ac regions | 0.08 | 1.11 | 0.37 | 2 | 0.74 |

Many of these annotation categories are highly dependent (Fig. 3), raising the question of what constitutes an appropriate correction for multiple comparisons. To estimate this correction we generated 10,000 simulated datasets of annotated mutations and assessed the correlation of p-values for the 51,801 categories across the simulations. Excluding categories with too few mutations to achieve nominal significance left 14,789 categories, and eigenvalue decomposition was used to estimate 4,211 effective tests based on the sum of eigenvalues that explain 99% of variation (Fig. 3). Correcting for 4,211 tests sets a category-wide significance threshold of 1.2×10^−5^ (Fig. 2a).

**Figure 3.**
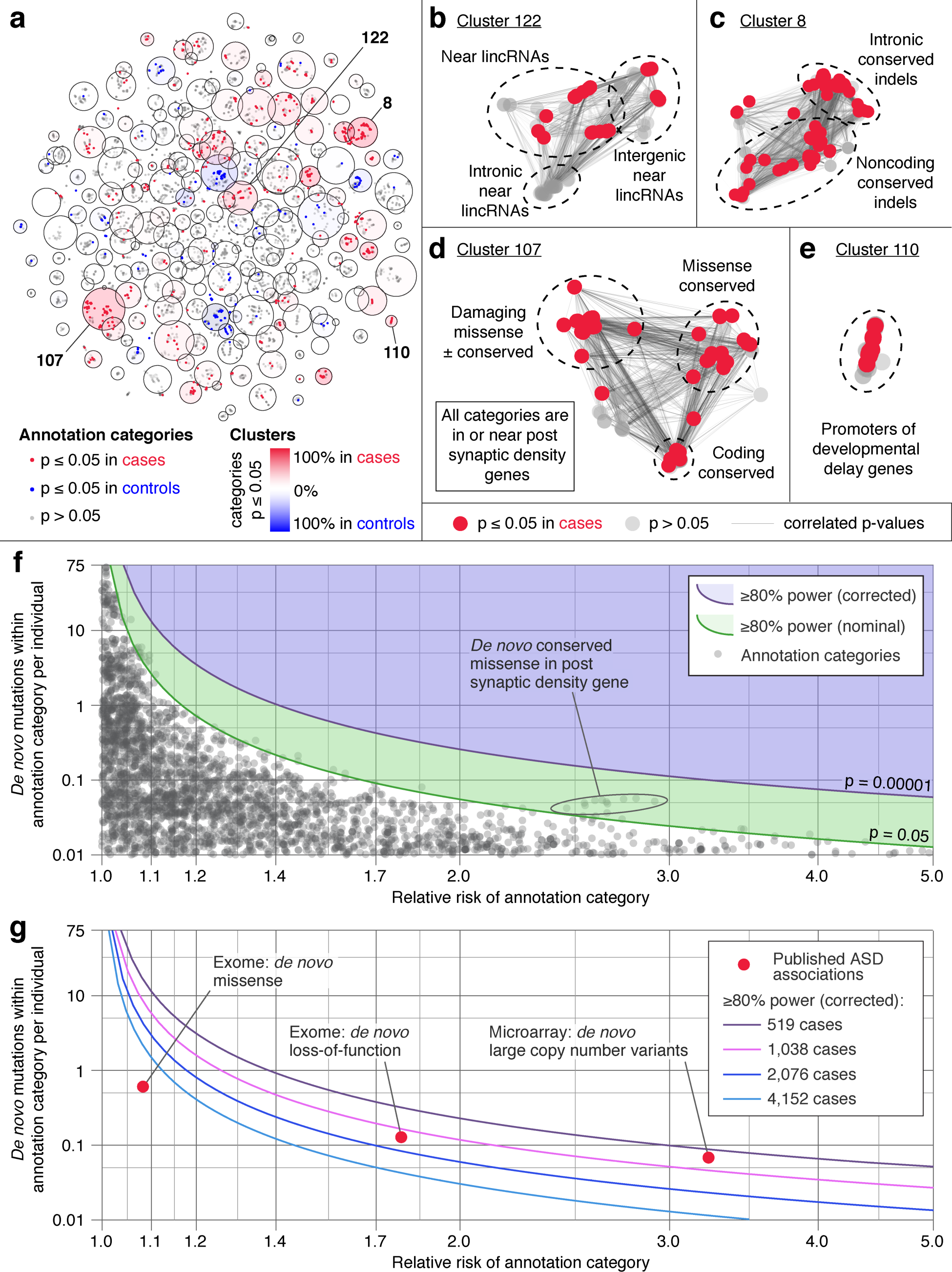
Effective number of tests in CWAS and power calculation. **a)** Correlations between p-values for 51,801 annotation categories across 10,000 simulated data sets were analyzed using Eigenvalue decomposition. After excluding tests with fewer than 7 variants in at least 50% of simulations, 14,789 categories remained; these are shown as a small dot with X and Y coordinates determined by t-Distributed Stochastic Neighbor Embedding. Red dots indicate categories that are nominally significant in cases, blue dots are nominally significant in controls, and grey transparent dots are not significant. Two hundred clusters of annotation categories were identified using k-means clustering and are represented as large circles with size determined by the number of effective tests required to account for the categories within the cluster. In total, 4,211 effective tests explain 99% of the variability in p-values. Clusters are colored according to the percent of nominally significant categories in cases (red) or controls (blue). Zoomed in plots from ‘a’ with edges representing p-value correlation are shown for: **b)** cluster 122, with 132 categories related to variants near lincRNAs that account for 41 effective tests; **c**), cluster 8, with 115 categories related to conserved indels that account for 30 effective tests; **d)** cluster 107, with 167 categories relating to variants in proximity to post synaptic density genes that account for 31 effective tests; and **e)** cluster 110, with 37 categories relating to promoters of developmental delay genes that account for 7 effective tests. **f)** The red line shows the threshold to achieve 80% power at nominal significance across the range of relative risks of a category (log_10_ scaled x-axis) and number of de novo mutations per individual within the category (log_10_ scaled y-axis). The blue line shows the 80% power corrected for 4,211 effective tests. The grey dots represent the observed results for de novo mutation burden in 519 families for the 11,876 annotation categories with ≥10 mutations. **g)** The lines show the threshold of 80% power across the range of relative risks and category sizes as sample size increases (correcting for correspondingly more effective tests). For reference, the results for well-defined categories of ASD risk are shown by the red dots.

While no single category met this threshold, we considered whether there was evidence of a tendency towards enrichment of categories in cases, suggesting an underlying signal. We therefore counted the number of nominally significant categories and compared this to expectation based on permutation and controls (Fig. 2b). We observed more significant tests than expected in cases in coding regions (p = 0.01) but not noncoding regions (p = 0.21), both overall and near ASD genes. This result gives important insight into genomic architecture; as cohort size increases we should anticipate that noncoding signal will remain weaker than the coding signal, unless annotation approaches improve dramatically. Moreover, since cases with known loss-of-function coding mutations were excluded from this sample, this suggests that the noncoding signal will likely be more modest than the signal from missense coding mutations. Interestingly, tests of annotation categories for *de novo* indels separate from SNVs showed a greater number of significant results than expected, and this enrichment was stronger for noncoding (p = 0.04) than coding indels (p = 0.10). Indels may represent a sweet spot for statistical power in interrogating the noncoding genome; they can disrupt regulatory elements to a greater degree than SNVs by virtue of their size while being detected in considerably greater numbers than SVs.

To further assess the role of rare noncoding variation for ASD we developed a polygenic risk score based on *de novo* variants, akin to similar scores developed previously for common and rare variants.^34,35^ The rate of *de novo* mutations in cases and controls was weighted based on the category RR and adjusted for p-value correlation structure (Fig. 3). Cross validation was used to select annotation categories that best predicted case-control status. In keeping with the modest differences observed between cases and controls, the derived score was not able to accurately predict case status, further supporting a limited role for rare noncoding mutations in this cohort. Of note, this model did not explicitly highlight the contribution of coding mutations, with the majority of selected categories relating to overall *de novo* burden (e. g. all variants, all intronic variants, and all intergenic variants). However, the model did highlight the role of two other functional annotations: conservation scores across vertebrate species and variants near long intergenic noncoding RNAs (lincRNAs, Fig. 3). Though neither finding is significant after correcting for multiple comparisons (Fig. 3), they present intriguing hypotheses for future studies.

Finally, we explored the impact of rare inherited SNVs and indels in the 405 families of European ancestry.^8^ Overall we observed a small excess of rare homozygous SNVs and indels (allele frequency <1%) in regions of homozygosity (ROH) in cases (66.1 per case vs. 63.1 per control; RR = 1.05; p = 2.4×10^−7^, one-sided binomial test). Since ROH blocks often contain multiple variants inherited simultaneously, we counted only one variant per ROH block and excluded variants in ROH blocks that overlapped coding regions. No significant excess of variants remained (3.53 per case vs. 3.51 per control; RR = 1.004; p = 0.91). No overall excess of rare heterozygous SNVs and indels was observed, including considering maternally and paternally inherited variants separately, and no category reached significance in a CWAS for either homozygous or heterozygous variants (Extended Data Figs. 11-21).

### Structural variation

Though no definitive noncoding signal was observed for small mutations, the strongest trends were observed in indels, in keeping with their larger size and presumed greater disruption to regulatory elements than SNVs (Fig. 2b). Following this logic, we assessed whether structural variants (SVs), which can rearrange and potentially disrupt large segments of the genome, might demonstrate a noncoding signal. We integrated the results of seven prediction algorithms to capture both changes in read-depth (three algorithms) and clusters of anomalously pairing reads indicating an SV breakpoint (four algorithms; see Online Methods). We then developed a series of *post hoc* algorithms, called RdTest, to correct for the limited concordance among individual algorithms (Extended Data Fig. 22). The method jointly tests for a significant difference in the read-depth signal supporting each predicted CNV against the normalized cohort background, and performs local k-means clustering to predict the likely presence of multiple copy states. We next integrated the statistically significant CNV segments with predicted balanced events using a series of breakpoint linking methods to identify signatures of 10 canonical balanced and complex SV classes,^36^ of which 64.5% altered copy number (e. g., paired-duplication inversion^37^) and 35.5% were copy number neutral.

These analyses identified a median of 4,089 SVs per individual, involving an average of 12.1 Mb of rearranged sequence per genome (Extended Data Fig. 23). Notably, these SVs result in a median of 84 loss-of-function and 21 whole-gene copy gain variants per person, and 7.5% of SV altered coding sequence compared to 2.2% of SNVs and indels. The variant frequency of SV in this cohort largely parallels that of SNVs and indels; 72.0% of all variants were rare (<1%) and 45.5% of variants were observed in only 1 family. In keeping with their presumed functional effect and resulting selective pressure, 61.4% of genic loss-of-function or copy gain SVs appeared in only a single family.

We compared standard WGS to 1,332 high quality CNVs previously reported from microarray data in the SFARI cohort (Extended Data Fig. 24),^1^ and observed an overall sensitivity of >99% and a 5.2% false discovery rate (FDR). We relied on long-insert WGS (liWGS; 3.5 kb inserts, median physical coverage of 102x) to validate SVs undetected with microarray (including small CNVs, copy-neutral balanced SV, and complex SV) and found a 4.3% overall FDR for 2,238 SV calls (Extended Data Fig. 24). Consistent with the comparisons to microarray and liWGS, cross-site validation using PCR and Sanger sequencing confirmed 92.3% of our predictions, suggesting high specificity from these analyses, very likely at the cost of sensitivity for small variants (see Methods), though we have no gold standard to determine this with certainty.

These analyses predicted 105 *de novo* SVs in the cohort, including 92 germline and 13 apparent mosaic SVs (Extended Data Figs. 25-27). In addition, we found that five subjects had sex chromosome aneuploidies (0.7% of SSC probands, 0.2% of siblings; Extended Data Fig. 28), and discovered nine SVs initially predicted to arise *de novo* that demonstrated evidence of germline mosaicism in a parent. Given the rarity of *de novo* SVs, there were limited data to derive insights comparable to those from *de novo* SNVs and indels. There was no significant difference in *de novo* SV burden between cases and controls (see Methods for sibling comparisons), though we did observe a small increase in risk among cases (RR = 1.53, p = 0.07). There was also a non-significant enrichment in ASD cases for *de novo* SVs localized to exons (2.3% versus 0.6%; RR = 3.7; p = 0.06), suggesting that there is a slightly increased burden of previously undetected SVs that disrupted protein coding sequence in ASD cases, and this result was more pronounced if we excluded multi-allelic (n = 20) and mosaic (n = 13) SVs (RR = 9, p = 0.02). There were *de novo* SVs that represented potential loss-of-function variants within ASD-associated genes, which included an exonic deletion of *CHD2* and a balanced translocation that disrupted *GRIN2B* (Fig. 4). Several other genes were disrupted by SVs in cases that were predicted to be intolerant to loss-of-function mutations (pLi ≥ 0.9^32^), but not associated with ASD from TADA analyses (*LNPEP, PAK7, SAE1, ZNF462, DMD*), while one such disruption occurred in a sibling (*USP34*). Overall, these analyses suggest that *de novo* loss-of-function SVs that were intractable to microarray may translate to a 1.7% increased burden in ASD cases compared to siblings, in addition to the 10.5% increased burden of cases harboring ASD relevant loss-of-function coding mutations and CNVs identified previously.^1^

**Figure 4.**
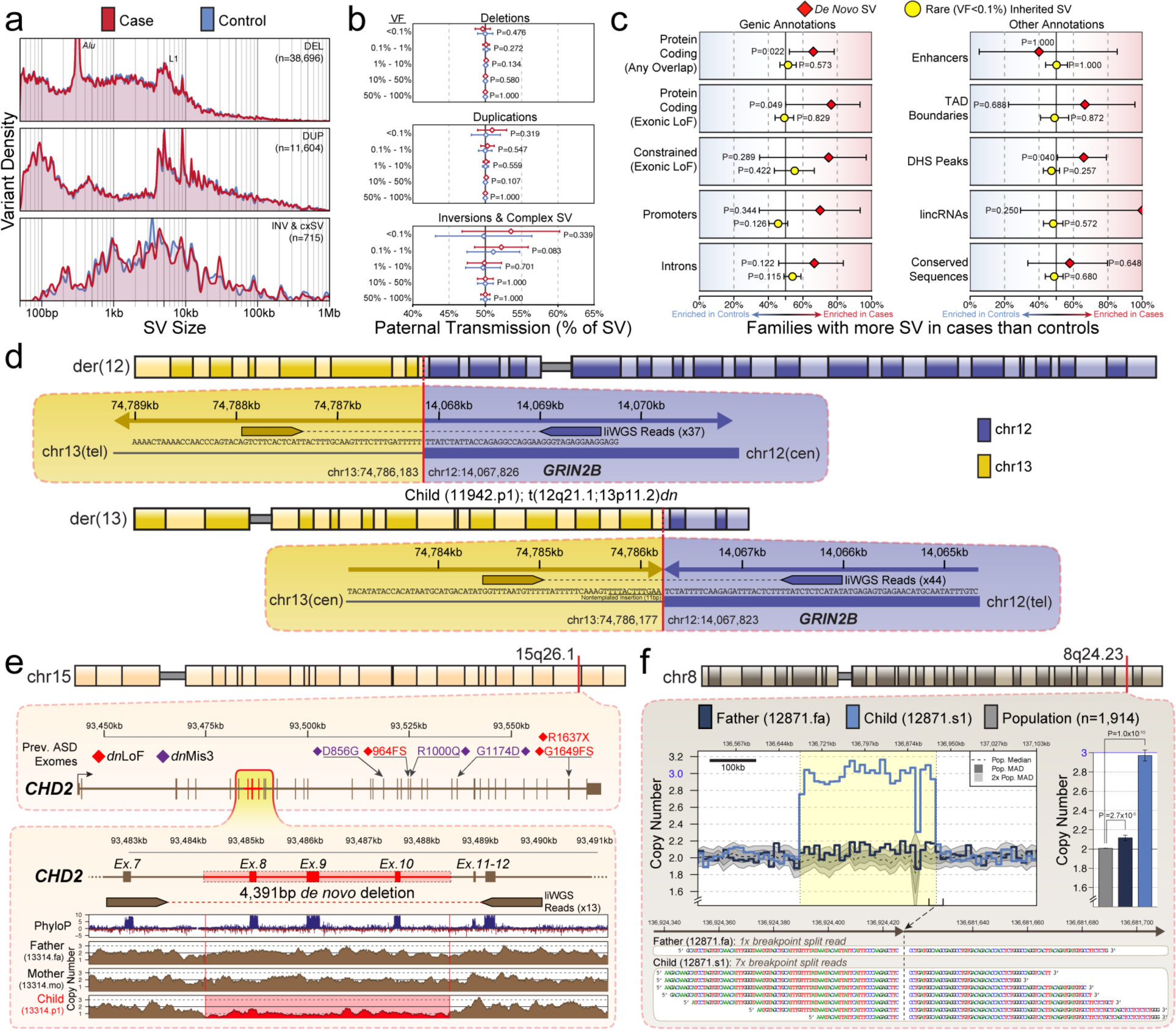
Structural variation in 519 ASD families. Structural variation (SV) analyses identified an average of 4,096 SVs per genome and 105 de novo SVs in this cohort. **a)** From analyses of these variants we observed no difference in distribution of SV sizes between cases and sibling controls for any class of SV (cxSV = complex SV). **b)** The majority of inversion variation detected in these samples (64.8%; 463/715) were complex, non-canonical rearrangements that fit previously described subclasses of complex SV.^36^ **c)** We observed no significant enrichments for either de novo or rare inherited SV (variant frequencies [VF] < 0.1%) in genic or noncoding annotations in cases versus controls after correcting for multiple comparisons. **d)** Analysis of balanced SV discovered a de novo reciprocal translocation in a case predicted to disrupt GRIN2B, a constrained gene previously implicated in ASD by recurrent de novo mutations.^1,32^ **e)** WGS revealed thousands of small CNVs undetected by previous analyses, including a 4,391bp de novo deletion of exons 8-10 of CHD2, a gene implicated in ASD due to recurrent de novo loss-of-function and missense point mutations from whole-exome sequencing.^1^ **f)** Analysis of breakpoint sequences also classified 13 de novo SVs that were predicted to be germline mosaic in the parents, such as a 364.2kb paternally transmitted mosaic duplication at 8q24.23 that was previously characterized as de novo in the child.

We next explored the properties and potential impact of 16,906 rare inherited SVs in the SSC (MAF <0.1%). Consistent with our previous analyses of large SV in the SSC,^36^ rare SVs were enriched for many of the hallmarks of selection against deleterious variation in the human genome when compared to common SVs (MAF > 1%), as they were more likely to disrupt genes (p=1.17×10^−81^), particularly constrained genes (p=7.00×10^−14^), and enhancers obtained from Fantom5^38^ samples (p=3.40×10^−51^). However, there was no significant difference between ASD cases and controls in the predicted impact of rare inherited SVs in this study, including no difference in overall size, percent of genome rearranged, or distribution of complex SVs (Fig. 4; Extended Data Fig. 23). We also did not detect any changes in SV burden in proximity to genes, or any signal when surveying up to 1 Mb from the transcription start site of genes. This result remained negative when we restricted analyses to variants in close proximity (2 kb) to constrained genes (min p = 0.25) and ASD-associated genes (min p = 0.69). The strongest noncoding signal in a CWAS analysis of SV was an increased burden of rare inherited SV (MAF < 0.1%) within introns of constrained genes (p = 0.0008), though this result was not significant when correcting for the considerable number of tests performed (see effective tests above). Finally, we identified signatures of large SVs that were not detected by microarray in the SSC, revealing that 0.9% of ASD cases (N=5) harbored a large balanced chromosomal abnormality (>3 Mb), and 429 CNVs >40 kb were detected by WGS but not microarray (Extended Data Fig. 24). Despite this improved power and resolution for SV detection, we found no significant differences in the rate of rare inherited SV as a mutational class in ASD, nor did we observe any evidence of biased transmission of any class of SV from either parent (Extended Data Fig. 29).

### Prediction of biologically relevant noncoding loci

The analyses reported above took an unbiased approach to testing the association of noncoding variation with ASD and it thus required appropriate correction for the effective number of tests performed. One could argue that, while we don’t have the same triplet code in the regulatory genome, there is good evidence to define one or more putative functional loci *a priori* that influence risk. Indeed, members of our consortium performed such analyses in the initial 39 pilot quartets, leveraging prior discovery of convergent co-expression of ASD genes in the midfetal prefrontal cortex^39^ to identify noncoding target regions as a single hypothesis (unpublished analysis). To define these regulatory targets, they generated H3K27ac ChIP-Seq data to identify regions of active transcription from 4 *post mortem* human brains (15-22 weeks post-conception, prefrontal cortex) and ATAC-Seq data to identify regions of open chromatin from 5 brains (16-22 weeks post-conception, prefrontal cortex). Previously published analyses have also suggested associations with noncoding regulatory variation through targeted biological hypotheses. These include association with variants localized to fetal CNS DNase I hypersensitive sites (DHS) within 50 kb of ASD-associated genes among these 39 SSC pilot quartets and 14 additional families,^25^ as well as a recent report of paternally inherited SV predicted to disrupt fetal brain promoters or UTRs of constrained genes in a study that included these SSC quartets.^40^

Despite the strong evidence for biological relevance in our unpublished pilot analyses, and an *a priori* association in a subset of these same families, our targeted hypothesis was refuted in the larger cohort: there was no excess of *de novo* mutations within these regions of open or active chromatin in the midfetal human prefrontal cortex (Extended Data Fig. 30). Similarly, no excess of mutations was observed by further filtering to variants in proximity to 179 ASD genes defined by WES at a false discovery rate of 0.3^1^ (Extended Data Fig. 30-31). Contrary to previously published analyses, we also find no evidence of enrichment for disruption of DHS sites in proximity to all genes, or ASD-associated genes, at any sliding window distance extending up to 1 Mb (Extended Data Fig. 32), nor did we observe enrichment of paternally inherited SV disrupting any class of functional annotation in proximity to all genes, constrained genes, or those genes previously associated with ASD.

### Integration and estimation of noncoding risk in ASD

An excess of *de novo* loss-of-function mutations and of *de novo* missense mutations has previously been described in WES data with RRs of 1.75 and 1.15, respectively.^14^ Resampling these WES data finds that about 300 families are required to observe the *de novo* loss-of-function burden (80% power, alpha = 0.05), while over 1,500 families would be necessary to observe the *de novo* missense burden (Fig. 3). If we count the number of *de novo* missense mutations in cases versus controls in the current WGS sample, the RR is only slightly inflated in cases (414/404 = 1.02) and it is not significantly different than 1.00, as expected from this power calculation. If, with the benefit of hindsight, we consider only 179 genes previously associated with ASD at a liberal false discovery rate of 0.3^1^ as a sole endpoint of our analyses, we find a much higher RR of 2.6 (21/8), which is significantly different from 1.0 (p = 0.01, one-sided binomial test, Fig. 2). As noted, however, this result does not survive correction for multiple comparisons and it is probably somewhat biased by the inclusion of these 519 families in the original WES analyses that defined the 179 genes. Moreover, filtering missense mutations instead by conservation, constrained genes, or brain-expressed genes, does not yield nominally significant evidence for risk.

These results give important context to interpreting the WGS data for 519 families and for the larger sample sets of the future. At 519 families, we should expect a noncoding signal equivalent to *de novo* loss-of-function to be nominally significant (alpha = 0.05), but not expect this of a signal equivalent to *de novo* missense until the sample size exceeds 1,500 families. As noted (Fig. 2), the noncoding signal we observe is weaker than that seen for *de novo* missense mutations. Furthermore, the best chance of achieving a significant test lies in integrating data that enriches for ASD-associated signal, such as proximity to ASD genes. Yet, when we searched over the space of *de novo* SNVs, indels, SVs, and rare homozygous variants, they showed no detectable concentration near *bona fide* or even likely ASD genes. Nor did these variants concentrate in any particular region of the genome, as could occur if disruption of a particular noncoding region were associated with large relative risk. Finally, they did not concentrate notably in any annotation category that we tested.

Without the triplet genetic code of the protein coding sequence we could not have distinguished loss-of-function, missense, and silent variants in the exome data and would expect a RR of 1.12 for all *de novo* mutations in coding regions. We would require 1,000 families to detect this burden (80% power, alpha = 0.05), over three-fold more than required to detect loss-of-function alone. This analogy represents the challenge of assessing noncoding regulatory risk from WGS data, exacerbated by the likelihood that regulatory variants are, as a group, unlikely to confer the same level of risk as loss-of-function variation. Moreover, because we have yet to discover the functional elements critical for disease risk, rather than specify them *a priori*, it induces a search over a large number of putatively functional elements and mandates far more stringent thresholds for statistical association as we have used.

To estimate the sample sizes required to discover annotation categories enriched for noncoding variation, we performed a power calculation across estimates of RR and numbers of variants per annotation category. Because these categories show complex correlation structure, and therefore simple corrections for multiple testing are inappropriate, we used eigenvector analysis to estimate the effective number of tests conducted. As sample size increases, the correction for number of categories becomes somewhat larger due to increased likelihood of observing a total number of *de novo* mutations in any given annotation category that is sufficient to achieve significance: the number of effective tests increases from ≈4,200 at 519 families to ≈7,600 at 4,000 families and approaches an asymptote of ≈10,000 (Fig. 3). The multiple testing burden produces a threshold for statistical significance on the order of 5 × 10^−6^. In this setting, over 4,000 families would be necessary to discover a noncoding element equivalent to missense variation.

## Conclusion

Refinements in DNA sequencing, computing capability, and statistical analyses now permit simultaneous evaluation of the coding and noncoding genome in many thousands of individuals. This eventually will precipitate a sea change in how we interpret the impact on ASD risk of rare variation throughout the genome. Yet, the complexity of the noncoding genome complicates interpretation for both *de novo* and inherited variation, and there are perils in underestimating its complexity. *A priori* prediction by experts of which regulatory elements of the noncoding genome should be important will limit the number of tests evaluated, and one could argue this limits the required correction for multiple testing. We find this argument wanting in terms of establishing a robust, unbiased framework to interpret disease association. Perhaps the simplest way to understand why is by analogy to common variants and a comparison of current-day genome-wide association studies (GWAS) versus the candidate gene tests of a previous era. GWAS results have a good record for replication, in large part because the field requires, for any study, large samples and appropriate correction for multiple testing. By contrast, despite investigator intuition about what genes are important to disease risk, candidate gene studies have had a miserable record regarding replication. This history of candidate gene studies, with a plethora of false positive and a paucity of true results,^41^ should make us highly skeptical of methods based on investigator-selected *a priori* hypotheses in the noncoding genome. Continuing the analogy, instead of candidate genes, the field would be substituting “candidate annotations”, with all likelihood of worse outcomes, due to myriad combinations of annotation, cell type, brain region, and developmental stage.

We anticipate that large-scale functional assays will continue to provide increasingly insightful annotation of the regulatory genome enabling future studies to better characterize and quantify the precise contribution of noncoding regulatory variation to ASD. In addition, high-throughput methods to validate noncoding variant function, such as STARR-Seq,^42^ for which there is no equivalent for coding missense mutations, could refine noncoding signals, potentially to the degree of implicating specific noncoding loci. Until that time, we recommend the GWAS path for WGS studies: rigorous evaluation of multiple hypotheses and appropriate correction for that multiplicity, as we have outlined here. If we hold to these standards, it will require very large sample sizes to make headway, but we predict that the ensuing inferences will be sound and replicable.

## METHODS

### Sample selection

519 quartet families (2,076 samples) were selected from the Simons Simplex Collection (SSC). The families were selected on the basis of having no known *de novo* rare CNVs, *de novo* loss-of-function mutations, or inherited rare CNVs at known ASD loci in the proband. The first 39 families were additionally selected for high paternal age, low IQ, and female sex while the second 480 were selected at random from the SSC. All of the families had pre-existing microarray data^1^ and pre-existing WES (47 trios without a sibling and 472 quartets)^14^. A complete list of the 2,076 samples is shown in.

### Whole genome sequencing

Whole blood-derived DNA from all four family members was transferred from the Rutgers University Cell and DNA Repository (RUCDR) to the New York Genome Center (NYGC). Rigorous quality control for the DNA led to 21 families being excluded prior to sequencing. The remaining 519 families were submitted for WGS. Data for the first 39 families was generated using PCR-based library preparation followed by sequencing on an Illumina Hi-Seq 2000. The next batch of 480 families were sequenced by PCR-free library preparation on an Illumina Hi-Seq X Ten. Sequencing reads for all samples were 150 bp paired-end cycles with a median insert of 423 bp. Sequencing yielded a median alignment rate of 99.3%, a strand balance of 0.50, a 0.11% duplication rate, and a median coverage of 37.8X per individual.

### Data processing

Using the NYGC processing pipeline, FASTQ reads were aligned to the hg19 reference from the 1000 Genomes Project (GRCh37.63) using BWA-mem version 0.7.8-r455. Reads were sorted and duplicates were removed with Picard, version 1.83. Indel realignment, base quality score recalibration, and variant calling with the GATK haplotype caller were performed using GATK version 3.1-1-g07a4bf8 for 19 families of the first batch, version 3.2-2-gec30ce for 21 families of the first batch, and version 3.4-0-g7e26428 for all 479 families of the second batch.

The BAM and gVCF files for 519 quartet families (2,076 samples) were transferred to Amazon Web Services (AWS) S3 storage system where they are available to access and download. For downstream steps on AWS, we deployed the CfnCluster based on the Lustre cluster system and multiple m4.10xlarge instances (amazon AMI: ami-3a081f50). We used GATK version 3.4-46-gbc02625 and the protocol detailed by GATK best practices (https://software.broadinstitute.org/gatk/best-practices/) to merge individual gVCF files into a combined VCF file. SNP and indel recalibration was then run on this combined VCF file. Variant Quality Score Recalibration (VQSR) metrics were created from a training set of highly validated variant resources: dbSNP build 138, HapMap 3.3, 1000 Genomes OMNI 2.5, and 1000 Genomes Phase 1. For the following analysis, we excluded variant calls with VQSR tranche level between 99.9 and 100%, and variant calls located in low-complexity regions^43^, as these calls have a high error rate or unusual characteristics^43,44^.

For annotation and subsequent analyses, indels were realigned using left-normalization, and multiple variants at the same locus were split into individual VCF lines using BCFtools. VCFs for each of the 519 families were then extracted from the combined VCF using BCFtools^45^, while retaining allele frequency and count information calculated from the full cohort. Spanning deletions were excluded from the family VCFs using a custom python script.

### Genomic prediction of common variant contribution in SSC cohort

Microarray data were limited to samples from the main European ancestry (1,634 families). We used a jackknife approach to determine genomic prediction within the SSC proband and pseudo-control samples: for each step of the jackknife, a proband and pseudo-control (comprised of the un-transmitted SNP alleles from mother and father) from one family was removed from the data^46^. Solutions on the observed (0/1) scale for the remaining individuals were obtained using mixed linear model equations taking into account 7 ancestry eigenvectors based on the genetic ancestry of probands and pseudo-controls. Heritability for ASD on the liability scale was 0.396, which was transformed to a heritability on the observed scale of 0.718 based on a prevalence of 0.01 and a 50: 50 ratio of cases and controls in our sample. Genomic predictions for the two samples left out were based on the linear regression of the known solutions using the genomic relationship matrix among probands and pseudo-controls from all families. Genomic predictions were scaled to have mean 0 and standard deviation 1. The SSC sample was divided into three groups, WGS sample (N=519, only 327 met our strict criterion for European ancestry), cases carrying damaging *de novo* mutations (N=438), and neither (N=869). Next we conducted an analysis of variance to determine if the mean genomic scores for the three groups were significantly different (in statistical package R, function ‘aov’).

### Variant annotation

Variants were annotated using Annovar^47^ and Bamotate^1^ in five groups:

1. *Variant type:* SNVs and indels were obtained from the final VCF and subject to the ROC-based filtering for high-quality variants. Indels are limited to the size less than 50 bp. SVs include deletions, duplications, insertions, and complex events.
2. *Gene-defined annotation:* Gencode complete version 19 (wgEncodeGencodeCompV19)^48^ gene definitions were obtained from the UCSC table browser (https://genome.ucsc.edu/). Variants were annotated against these gene definitions using Bamotate; where multiple possible annotations were present they were assigned in the following order of priority: coding, intron, promoter, UTRs and intergenic. Promoters were defined as 1kb upstream of the transcription start site (TSS). For intergenic variants the nearest TSS was also identified.
3. *Annotation of species conservation scores:* To evaluate the conservation status of identified variants, we used two conservation metrics: phastCons 46-way scores, and phyloP scores from a 46-way vertebrate comparison from the UCSC table browser^49^,^50^.
4. *Annotation of gene sets:* Gene lists were chosen based on prior association with ASD (e. g. post-synaptic density genes). ASD risk genes (FDR<0.3) were obtained from Sanders et al. (2015)^1^. Genes co-expressed with ASD genes were defined as the union of the two co-expression modules identified by Willsey et al. (2013)^39^ in the: 1) human midfetal prefrontal and primary motor-somatosensory cortex; and 2) infant mediodorsal thalamic nucleus and the cerebellar cortex. Genes associated with developmental delay were downloaded from the Development Disorder Genotype - Phenotype Database (https://decipher.sanger.ac.uk/ddd)^26,51^ in Sept 2016. The 2,156 genes were filtered to: 1) confirmed DD gene; 2) predicted as loss-of-function in the mutation consequence; and 3) including term “Brain” in the organ specificity list. CHD8 target genes were defined as the union of lists from two previous ChIP-Seq studies^52,53^, and FMRP target genes were selected from Darnell et al. (2011)^33^. Human cortex post-synaptic density (PSD) proteins were downloaded from the Genes2Cognition database (http://www.genes2cognition.org/)^54^. Constrained genes were defined as probability of being loss-of-function intolerant (pLI) score≥0.9 in the ExAC database^32^. If a variant was within a Gencode transcript then that transcript was cross-referenced to these gene lists. For intergenic variants, the nearest transcription start site was cross-referenced to these gene lists.
5. *Annotation of regulatory regions:* BED files were obtained for multiple regulatory regions. Known enhancers were downloaded from the Vista enhancer annotation (vistaEnhancers) from the UCSC genome browser^55^ and the pre-defined enhancer set from the FANTOM 5 server (http://enhancer.binf.ku.dk/presets/)^38^. ENCODE-defined transcription factor binding sites and DNase hypersensitive sites were downloaded from UCSC genome browser (wgEncodeRegTfbsClusteredV2 and wgEncodeRegDnaseClusteredV3). Human accelerated regions (HARs) were obtained from Doan et al. 2016^56^.

For histone marks and chromatin states, we utilized data from the NIH Roadmap Epigenome Project^57^. For histone marks and chromatin states, we merged data from brain tissues (E067 Angular Gyrus E068, Anterior Caudate, E069 Cingulate Gyrus, E070 Germinal Matrix, E071 Hippocampus Middle, E072 Inferior Temporal Lobe, E073 Mid Frontal Lobe, E074 Substantia Nigra, E081 Fetal Brain Male, E082 Fetal Brain Female), neurospheres (E053 neurosphere cultured cells cortex derived, E054 neurosphere cultured cells ganglionic eminence derived), ES-derived neuronal cells (E007 H1-derived neuronal progenitor cultured cells, E009 H9-derived neuronal progenitor cultured cells, E010 H9-derived neuron cultured cells), and astrocytes (E125 NH-A Astrocytes).

In addition to the Roadmap Epigenome Project and ENCODE data, we utilized data sets generated at UCSF from mid-fetal human prefrontal cortex tissue (15-22 gestational weeks). These data sets included ATAC-seq, to identify regions of open chromatin, and ChIP-seq for H3K27ac, to identify putative active enhancer regions. Peaks were called by MACS (H3K27ac ChIP-seq) and Homer (ATAC-seq). Identified peaks common to two or more individual samples (1≥bp overlap) were used for annotation.

### Detection of high quality SNVs and indels

As we had no established best practices or predetermined filtering criteria available for rare variants in WGS data, we developed an optimized set of thresholds for various quality metrics to detect rare SNVs and indels. For this, we compared two sets of rare variants which have the most distinct quality metrics – 1) private transmitted variants (only observed in one family and no frequency given in the 1000 Genome Project or ExAC database), which are likely true variants, and 2) variants that are Mendelian violations in at least one child but are also observed in an unrelated individual, which are likely false positive calls. The ability of individual quality metrics obtained from the final VCFs to distinguish these true variants from false variants was assessed using receiver operating characteristic (ROC) curves. The metric and threshold that yielded the maximum increase of specificity and the minimum decrease of sensitivity was selected after which the training set was filtered by these criteria and the process repeated. This sequential ROC analysis was repeated until we no longer observed improvement in sensitivity and specificity.

### Detection of high quality *de novo* SNVs and indels

Four algorithms run on the default settings were used to detect *de novo* SNVs, TrioDeNovo^58^, DenovoGear^59^, PlinkSeq (https://atgu.mgh.harvard.edu/plinkseq/), and DenovoFlow. For *de novo* indels, DenovoGear was replaced with Scalpel^60^. DeNovoFlow is a custom script that parses all possible Mendelian violations from each family, given GATK quality metrics. The union of these four algorithms made predictions for 86,921 Mendelian violation SNVs and 5,726 indels per child.

These numbers are large, suggesting a high false positive rate among putative *de novo* calls. To identify high quality *de novo* variants from the call set, we applied the same sequential ROC approach as above with true positive calls defined by PCR Sanger validation *de novo* mutations from prior work (1,302 selected SNVs; 95 selected indels). Sequential ROC curve analyses were applied to all variant- and individual-level quality metrics for the child and both parents. This analysis predicted 87.3% sensitivity and 98.8% specificity for SNVs using 3 additional metrics, and 86.3% sensitivity and 93.0% specificity for indels using 4 additional metrics.

### Validation of high quality *de novo* SNVs

From the 66,366 high quality *de novo* SNVs, 250 mutations were selected at random (based on available DNA) for validation in the child and both parents using PCR amplification and high-throughput sequencing on an Illumina MiSeq. We examined PCR products from all 250 child reactions on a gel and 13 (5%) failed to make a product and were excluded from the analysis. Of the remaining 237 putative mutations, we observed an overall mean coverage of 26,818X. Based on investigation of off-target coverage, we determined that a depth coverage ≥ 50X was required to ensure an accurate genotype and any samples that failed to achieve this coverage were considered sequencing failures due to insufficient depth. All putative mutations in the child met this threshold, however for 7 of these, no variant was detected in the child. In the remaining 230 putative mutations, 18 had insufficient coverage in one or more parents and were excluded from the analysis. The remaining 212 putative mutations with sufficient coverage in the child and both parents all validated as *de novo;* no inherited variants were observed. Our overall confirmation rate for *de novo* SNVs was therefore 96.8% (212/219; 212 validated versus 7 with sufficient coverage but no variant in the child).

### Validation of high quality *de novo* indels

From the 9,961 high quality *de novo* indels, 250 indels (125 non-coding deletions and 125 non-coding insertions) were selected at random for validation using PCR amplification and high-throughput sequencing on an Illumina MiSeq. Of these, 16 were larger than 50bp and were excluded from the analysis (*de novo* confirmation rate of 6%). We examined PCR products from all of the remaining 234 child reactions on a gel and 7 (3%) failed to make a product and were excluded from the analysis. Of the remaining 227 putative mutations, we observed an overall mean coverage of 19,461X, however 7 failed to meet our threshold of ≥ 50x coverage in the child and were excluded from the analysis. Of the remaining 220 putative mutations, 75 failed to identify a variant in the child despite adequate coverage. In the remaining 145 putative mutations, 8 had insufficient coverage in one or more parents and were excluded from the analysis. Of the remaining 137 putative mutations with sufficient coverage in the child and both parents, 131 validated as *de novo* and 6 were inherited from one parent. Our overall confirmation rate for our first round of *de novo* indels <50bp was therefore 61.8% (131/212; 131 validated versus 6 inherited indels and 75 with sufficient coverage but no variant in the child).

Based on the results of this first round of validations, *de novo* indel prediction was refined identifying 5,932 mutations overall, and a second round of validation was performed on 200 randomly selected variants <50bp. From this final validation set, 189 (94.5%) putative mutations achieved adequate coverage in the child, but 28 of these failed to identify a variant in the child. Of the remaining 161 variants, 13 had insufficient coverage in the parents and were excluded from the analysis. In the remaining 148, 145 were validated as *de novo*, while 3 were inherited. Therefore, with the improved indel filtering criteria, 82.4% of putative mutations were confirmed as *de novo* (145/176; 145 validated versus 3 inherited indels and 28 with sufficient coverage but no variant in the child), showing a significant improvement relative to the exploratory analyses.

### Validation of mutations in ASD-associated genes

We also attempted validation for four putative mutations in known ASD-associated genes: one SNV in *ADNP*, chr20: 49548007; two SNVs in *GABRB3*, chr15: 26327365 and chr15: 26327513; and one indel in *NRXN1*, chr2: 51259257. All four mutations were validated as *de novo*.

### Detection of high quality *de novo* structural variants

#### Algorithm integration and variant adjudication

We used a two-tier SV detection pipeline, in which we integrated four paired-end/split-read (PE/SR) algorithms and three read-depth (RD) algorithms to discover a maximal list of candidate SV loci, then adjudicated each predicted variant with a joint analysis of the cohort that included a statistical test for likely *de novo* status of each alteration. Our pipeline incorporated PE and SR calls from Delly v0.7.3,^61^ Lumpy v0.2.13,^62^ Manta v. 0.29.6,^63^ and WHAM-GRAPHENING v1.7.0,^64^ each of which was run jointly on the four members of each quad. We included read-depth calls from GenomeSTRiP v2.00.1696,^65^ CNVnator v0.3.2^66^, and cn. MOPS v1.8.9^36^. We developed a read depth verification algorithm in R (RdTest) to determine the likelihood of true dosage alterations at a candidate locus by testing for statistically significance differences in depth between samples with disparate copy states. The detection of SV in repetitive regions of the genome remains challenging,^67^ as variant prediction in these regions frequently relies only on depth evidence. While remaining cognizant that many CNVs in the human genome are mediated by such repeats, we sought to prioritize specificity over sensitivity for SV calls within these regions and performed a series of ROC curve analyses to identify filters which would minimize the frequency of false positive variants produced in repetitive and low-complexity segments. From these analyses, we restricted SV predictions to exclude sites of multiallelic SV (k ≥ 6) and required any SV with only read-depth evidence to be at minimum 4 kb. We also performed the joint analysis of copy number difference in a batch-specific framework (pilot n=160 and Phase 1 n=1,916) to correct for the demonstrable differences in read-depth features between the datasets (which were PCR+ and PCR−, respectively), and further split the samples by sex for SV on allosomes. Notably, in adjudicating each variant, the metrics computed in the Phase 1 samples were used whenever available, and RdTest was also performed on a per-algorithm basis to filter spurious algorithm-specific calls. Finally, across all passing CNV we then genotyped homozygous deletions, defined as samples with a normalized read depth of less than 0.1 in at least half of the normalized read-depth bins. Notably, our analyses of sex chromosome SV revealed five samples with sex chromosome anomalies; three XXY Turner syndrome and two subjects with XYY syndrome (Jacob’s syndrome).

#### Distinguishing 10 classes of balanced and complex SV

In addition to our evaluation of polymorphic and *de novo* CNVs, we assessed the spectrum of balanced SV and complex SV in the SSC, as we have done previously in this cohort with large SVs.^36^ We applied the algorithm integration pipeline for PE/SR calls described above to obtain a set of candidate inversion and translocation breakpoints. We first used bedtools to overlap these breakpoints with the CNV loci predicted to be significant by RdTest to identify complex SV with large associated CNV, then to identify candidate pairs within the remaining breakpoints that could constitute a resolved SV. We resolved the variant structure at each of these loci by matching the ordering of breakpoints to complex SV signatures previously identified by Collins et al.,^36^ and used RdTest to evaluate read-depth support at novel CNV sites associated with complex inversions. We identified 19,342 observations of 127 such inversion-associated CNV between 300 bp and 4 kb that were not found with the CNV discovery pipeline, as they lacked canonical PE/SR evidence and were below RD-only algorithm resolution. In total, we identified 38,658 deletions, 11,598 duplications, 230 inversions, and 4 reciprocal translocations with this variant classification pipeline. Further, we discovered 453 complex SV across 8 classes, of which 99% included copy number alteration.

#### Validation of SV with microarray and jumping libraries

We compared the standard short-insert WGS (referred to as siWGS for clarity) SV calls to two previously published SSC datasets including long-insert WGS (liWGS, “jumping”) libraries on 456 of the 519 cases^36^ and microarray data available for all 2,071 samples with SV.^1^ To account for the differences in resolution across the three technologies, we restricted comparisons to variants which met three criteria: 1) a minimum size of 40 kb for microarray and 10 kb for liWGS; 2) at most 30% of the variant region localized to an annotated segmental duplication region, microsatellite, heterochromatin, or one of our defined multi-allelic regions; and 3) a variant frequency <10%. These filters were applied equivalently to the siWGS SVs in each comparison, resulting in 1,633 siWGS variants in the array comparison (Extended Data Fig. 24) and 2,238 siWGS variants assessed for support in the jumping libraries. Overall, we observed a 5.2% FDR based on the array data and a 4.3% FDR when comparing to the jumping libraries.

#### Validation of de novo structural variants

Validation was assessed on 68 *de novo* SV predictions using microarray, liWGS, and PCR followed by Sanger sequencing. PCR primers were designed using a custom script and Primer3,^68^ optimizing for sequencing data and the predicted size of the SV event. For one variant (DenovoCNV_53) in an AT-rich region we supplemented the validation with ddPCR. These initial exploratory analyses revealed CNV size (<700 bp) to be the predominant driver of false positive *de novo* predictions as our read-depth validation lacks sufficient data at this resolution, which led to a restriction on *de novo* SV predictions below this threshold to require support from two PE/SR algorithms in addition to RdTest adjudication. These methods returned a final validation estimate of 92.3% (48/52 test variants) with the final algorithm implementation.

#### SV annotation and statistical burden analyses

Each SV was annotated with any predicted overlap with the canonical transcript of 20,156 protein-coding genes in Gencode v19, as described above. In brief, deletions were considered loss-of-function (LoF) if they affected any coding sequence, duplications were considered LoF if they affected an exon but did not extend outside the transcript’s boundary, and inversions were considered LoF if one breakpoint localized to a coding exon or any genic space spanning the coding sequence (but not if the entire coding sequence was inverted). Duplications were considered to be “copy-gain” if they spanned the entirety of a transcript’s boundary. A variant was required to localize fully to an intron to be considered intronic, and each variant was additionally annotated with any gene whose UTR or promoter region (<1 kb upstream of TSS) it disrupted. These same criteria were applied to noncoding variation. Statistical burden testing was also performed using a CWAS design, paralleling the SNV analyses described above. Notably, families were selected after screening for probands harboring large and presumably loss-of-function *de novo* CNVs and coding mutations, but families with siblings harboring comparable mutations were not excluded. These analyses can impact estimates of SV association, and we consequently filtered any family in which the sibling met similar exclusionary criteria (n=27). We additionally excluded five families in which a family member demonstrated an aberrant WGS dosage profile that prohibited accurate SV prediction. Enrichment of rare SV was restricted to the 405 families with European ancestry described above in the SNV analyses.

## Acknowledgements

We are grateful to the families participating in the Simons Foundation Autism Research Initiative (SFARI) Simplex Collection (SSC). This work was supported by grants from the Simons Foundation for Autism Research Initiative (SFARI #385110 to N. S., A. J. W., M. W. S., S. J. S.; #385027 to M. E. T., J. D. B., B. D., M. J. D., X. H., and K. M. R.; #388196 to G. B., H. C., A. Q.; and #346042 to M. E. T.), the National Institute for Health/National Institute for Mental Health (R37MH057881 and U01MH111658 to B. D. and K. M. R.; HD081256 and GM061354 to M. E. T.; U01MH105575 to M. W. S.; U01MH111662 to M. W. S. and S. J. S. R01MH110928 and U01MH100239-03S1 to M. W. S., S. J. S, and A. J. W.; U01MH111661 to J. D. B.; U01MH100229 to M. J. D.), Autism Science Foundation to D. M. W., and the March of Dimes to M. E. T. We would like to thank the SSC principal investigators (A. L. Beaudet, R. Bernier, J. Constantino, E. H. Cook, Jr, E. Fombonne, D. Geschwind, D. E. Grice, A. Klin, D. H. Ledbetter, C. Lord, C. L. Martin, D. M. Martin, R. Maxim, J. Miles, O. Ousley, B. Peterson, J. Piggot, C. Saulnier, M. W. State, W. Stone, J. S. Sutcliffe, C. A. Walsh, and E. Wijsman) and the coordinators and staff at the SSC clinical sites; the SFARI staff, in particular N. Volfovsky; D. B. Goldstein for contributing to the experimental design; the Rutgers University Cell and DNA repository for accessing biomaterials; the New York Genome Center for generating the WGS data.

## Author Contributions

Experimental design, DMW, HB, JA, MRS, JTG, MJW, XH, NS, BMN, HC, AJW, JDB, MJD, MWS, AQ, GTM, KR, BD, MET, and SJS; Identified de novo SNVs and indels, DMW, JA, SD, MG, JDM, LS, AJW, and SJS; Identified structural variants, HB, JA, MRS, JTG, RLC, RML, AF, MG, REH, SK, LS, HZW, SAM, AQ, GTM, and MET; Confirmed de novo variants, DMW, SD, GBS, BBC, JD, CD, CE, HZW, and MJW; Annotation of functional regions, DMW, JA, SD, EM, JDM, YL, SP, JLR, NS, MET, and SJS; Generated midfetal H3K27ac and ATAC-Seq data, EM, TJN, ARK, and JLR; Developed genomic prediction score and de novo score, LZ, LK, KR, and BD; Analyzed SNVs and indels (Figs. 1 and 2), DMW, JA, and SJS; Analyzed SVs (Fig. 4), HB, MRS, JTG, and MET; Assessment of P-value correlations, effective number of tests, and power analysis (Fig. 3), DMW, JA, LZ, GBS, KR, BD, and SJS; Manuscript preparation, DMW, HB, JA, MRS, JTG, LZ, RLC, SD, BMN, HC, JDB, MJD, MWS, AQ, GTM, KR, BD, MET, and SJS.

